# Type I IFN Dynamics Define an Early Checkpoint for Survival and Orchestrate Systemic Neutrophil Heterogeneity in Lethal Viral Infection

**DOI:** 10.64898/2026.01.22.701212

**Authors:** Riho Saito, Akisawa Satomi, Hiroki Sugishita, Yukiko Gotoh, Tomohiko Okazaki

## Abstract

The early host response is critical for protection against viral infections, yet the systemic events that dictate individual differences in outcomes remain poorly defined. Here, we leveraged variability in survival following intranasal vesicular stomatitis virus (VSV) challenge in genetically identical mice to retrospectively profile systemic immune responses associated with survival or lethality. Survival was strongly associated with a robust systemic type I interferon (IFN) surge within 24 hours of infection, and blockade of type I IFN signaling during this narrow early window markedly reduced survival, establishing early IFN induction as a key determinant of outcome. This protective IFN surge rapidly remodeled the systemic immune landscape. Single-cell profiling revealed a transcriptionally and functionally distinct ICAM1⁺ neutrophil subset as the strongest early correlate of survival, primed within the bone marrow. ICAM1⁺ neutrophils exhibited a pro-inflammatory signature and enhanced phagocytic activity compared with ICAM1⁻ neutrophils, which predominated in lethal outcomes. Together, these findings define a type I IFN–driven early checkpoint that governs survival in lethal viral infection and identify ICAM1⁺ neutrophils as a blood-accessible biomarker of protective immunity, highlighting the critical role of early innate immune dynamics in shaping disease trajectories and providing a framework for early prognostic markers and host-directed therapies.

## INTRODUCTION

The severity of viral infections varies considerably among individuals, a phenomenon arising from the complex interplay among host, viral, and environmental factors ^1^ . Despite a growing understanding of risk factors, elucidating the precise mechanisms that determine the ultimate clinical outcome remains a major challenge. Addressing this challenge is critical for the early prediction of individual outcomes, thereby enabling timely therapeutic intervention.

Previous studies have tackled this challenge by comparing immune cell and cytokine profiles in patients with mild versus severe outcomes. These studies have provided important insights, revealing associations between myeloid dysregulation, lymphoid impairment, or aberrant cytokine responses and disease severity in infections such as SARS-CoV-2^2,3,4,5^ and influenza virus^6^. However, they are limited in their ability to elucidate the crucial early events of infection, as samples are typically collected only after symptoms have become apparent. Human challenge studies can probe the initial phase^7,8^ but are ethically restricted to inducing only mild outcomes, leaving the initial host responses that distinguish mild from severe outcomes largely uncharacterized.

Animal models are indispensable for dissecting these early, dynamic events. Intriguingly, significant variability in disease outcome often emerges even in highly controlled experimental settings where genetic and environmental factors are uniform. A classic example is the intranasal challenge of mice with vesicular stomatitis virus (VSV), which leads to a dichotomous outcome of either survival or death^9,10^.

The initial phase of a viral infection is governed by the immediate host response to pathogen sensing ^11, 12^ . This primary response, which precedes adaptive immunity, is critical as it shapes the entire subsequent disease trajectory. The efficacy of this innate response hinges on the rapid and robust induction of potent antiviral effector programs designed to control viral replication and orchestrate further immune involvement ^13,14^ . Much of the research on these early viral responses has focused on the local site of infection. In contrast, the early systemic response in the peripheral blood remains a key area of investigation, as it is a clinically accessible and ideal source for prognostic biomarkers. Therefore, identifying a predictive signature from blood at the earliest possible time point would be a major advance for timely therapeutic intervention. While some studies have characterized the early systemic immune response, a critical gap remains: few studies have directly linked cellular profiles from a remarkably early time point—such as 24 hours post-infection—to the ultimate survival outcome. This represents a major gap in our understanding of how early systemic responses shape the trajectory of a viral infection.

Type I interferons (IFNs) are universally recognized as a cornerstone of the early host defense^15^. Their rapid induction upon viral sensing triggers a wide range of interferon-stimulated genes (ISGs) that establish an antiviral state in host cells. While the necessity of type I IFN response for viral control is well-established, less is understood about the temporal dynamics of this response. The timing of type I IFN induction and the duration of its signaling can profoundly influence its effects, shifting the balance from a protective, virus-clearing response to a pathological, tissue-damaging one ^16, 17, 18^. However, the degree to which the precise timing of the initial type I IFN surge acts as a critical determinant of an individual’s ultimate outcome remains poorly defined.

Here, we propose that the variability in survival outcomes within genetically uniform mice under highly controlled conditions provides a unique opportunity to uncover the critical, early physiological differences that drive the divergence in infection outcomes, rather than being mere experimental noise. To explore the mechanisms underlying survival versus death following viral infection, we retrospectively analyzed immune cell profiles in the blood immediately before and shortly after infection. This revealed dynamic and transient changes in circulating immune cells that preceded the immune cell infiltration and activation at the infection site, which is previously described^19^. These early shifts were driven by a transient type I IFN response, which emerged as a key determinant of survival. Importantly, this robust, early type I IFN surge orchestrated a distinct systemic immune landscape, marked by the emergence of a functionally potent, pro-inflammatory neutrophil population characterized by high ICAM1 expression, which was more prominent in the survival group compared to the lethal group. Together, these findings establish a new framework in which the precise temporal dynamics of innate immunity, independent of host genetic variation, play a critical role in shaping the outcome of lethal viral infection.

## RESULTS

### Transient systemic immune dynamics in the peripheral blood during the early phase of VSV infection

While the intranasal vesicular stomatitis virus (VSV) infection model is known to induce progressive infiltration of immune cells into the brain beginning around 3 days post infection (dpi)^19^, the systemic immune dynamics in the peripheral blood remain largely uncharacterized. To investigate early changes in circulating immune cells, we comprehensively mapped the systemic immune landscape by performing single-cell RNA sequencing (scRNA-seq) on peripheral blood from mice at 0, 1, and 5 days post-intranasal VSV challenge (Fig.1A and Fig.S1A). This analysis revealed a transient wave of neutrophilia and lymphopenia at 1 dpi, characterized by a marked increase in circulating neutrophils and a parallel decrease in lymphocytes including B and T cells (Fig.1B, C, D), confirmed by flowcytometry analysis (Fig.S1B). By 5 dpi, these changes had largely resolved, returning to baseline levels (Fig.1B, C, D). Furthermore, a distinct population of interferon-stimulated gene (ISG)-expressing cells (ISG_high clusters) emerged in the blood at 1 dpi and disappeared by 5 dpi (Fig.1E). Together, these findings suggest that intranasal VSV infection triggers a rapid and transient remodeling of circulating immune cells, coupled with a systemic antiviral response during the early phase of infection.

**Figure 1.**
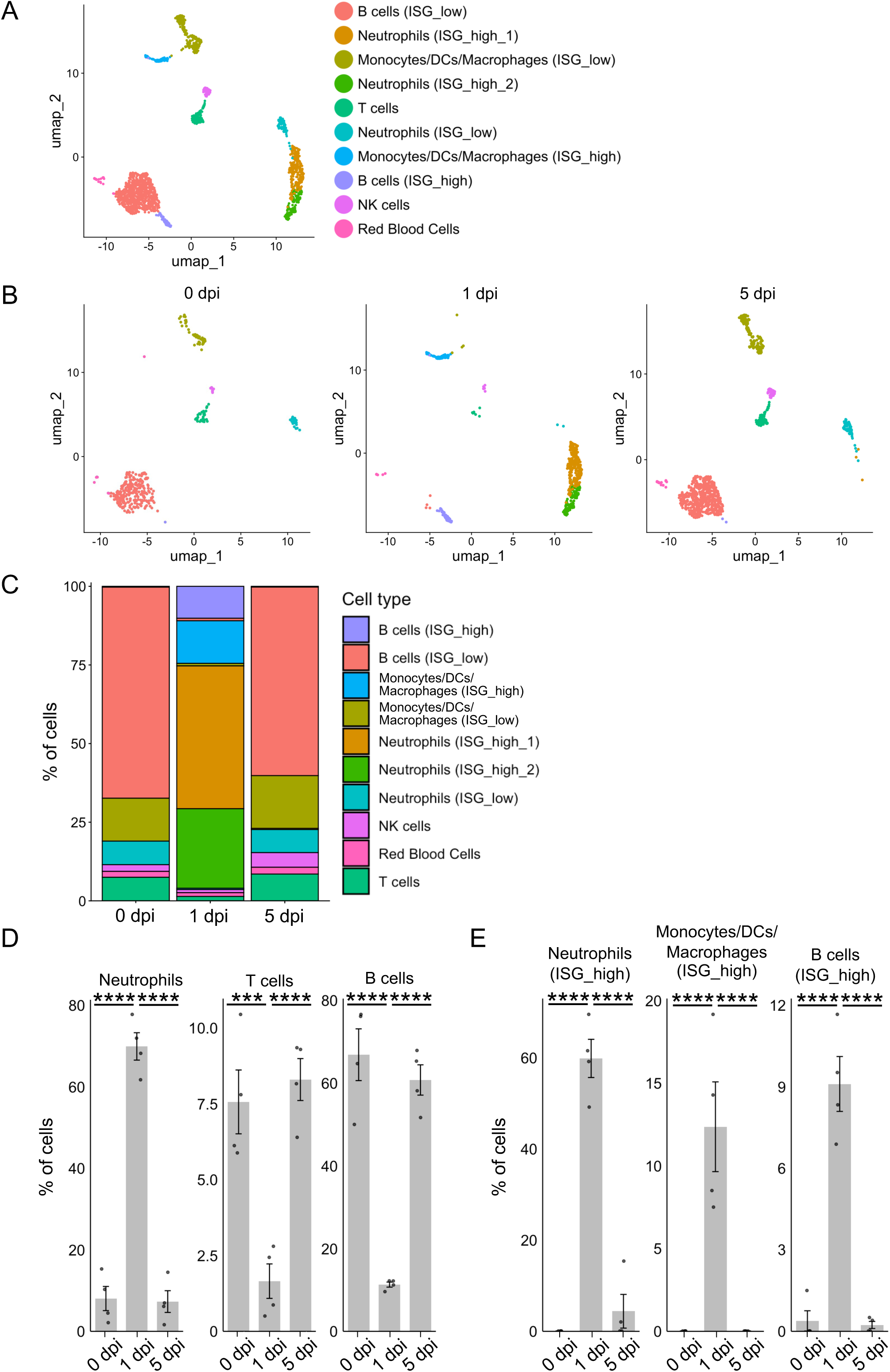
Transient systemic immune remodeling in peripheral blood during early VSV infection. **(A)** UMAP projection of circulating immune cells from the survival and lethal groups (n = 2 each) at 0, 1, and 5 days post-infection (dpi). **(B)** UMAP projections from (A) split by time point to show cells at 0, 1, or 5 dpi individually (cell numbers =374, 502 and 834 cells, respectively). **(C)** Percentages of each cell cluster within total circulating cells at each time point. **(D)** Percentages of neutrophils; Neutrophils (ISG_high_1) + Neutrophils (ISG_high_2) + Neutrophils (ISG_low), B cells; B cells (ISG_high) + B cells (ISG_low), and T cells clusters within total circulating cells at each time point. (n = 4 each) **(E)** Percentages of neutrophils (ISG_high); Neutrophils (ISG_high_1) + Neutrophils (ISG_high_2), Monocytes/DCs/Macrophages (ISG_high), and B cells (ISG_high) clusters within total circulating cells at each time point. (n = 4 each) Data in (D and E) are shown as mean ± SEM, where each point represents an individual mouse. \*\*\**P* < 0.005, **** *P* < 0.001. Statistical significance was determined using one-way ANOVA with Tukey’s post-hoc test (D and E).

### Transient interferon production in the draining lymph node contributes to systemic immune remodeling

Because our single-cell analysis revealed the transient emergence of an ISG-high population in circulating immune cells at 1 dpi, we hypothesized that this response might be driven by type I IFN production in the draining lymph nodes, as local viral infection is known to induce type I IFN not only at the site of infection but also in the draining nodes ^20^, thereby influencing systemic immune cell states^21^. We therefore examined the type I IFN induction in the superficial cervical lymph nodes (cLNs), which drain the nasal lymphatics and cerebrospinal fluid (CSF)^22^ and thus serve as the primary draining nodes in this infection model. Daily analysis from 0 to 4 dpi revealed a sharp and transient induction of IFN-α and IFN-β expression specifically at 1 dpi, accompanied by upregulation of multiple ISGs (Fig. 2A). In parallel, the expression of CD69—a key regulator of lymphocyte retention in lymph nodes by type I IFNs^23^ —was also transiently elevated at 1 dpi. Consistent with these results, robust accumulation of lymphocytes was observed in the cLNs but not in non-draining inguinal lymph nodes (iLNs) at 1 dpi (Fig. 2B), when circulating lymphocytes drastically decreased (Fig.S1B). Notably, type I IFN signaling blockade by the administration of an anti-IFNAR1 antibody at 0 dpi attenuated the transient lymphopenia, resulting in significantly higher numbers of circulating lymphocytes at 1 dpi compared to control (Fig. 2C). Collectively, these data indicate that the short-lived burst of the type I IFN production in the draining lymph node orchestrates the dynamic remodeling of circulating immune cells and the activation of systemic antiviral programs.

**Figure 2.**
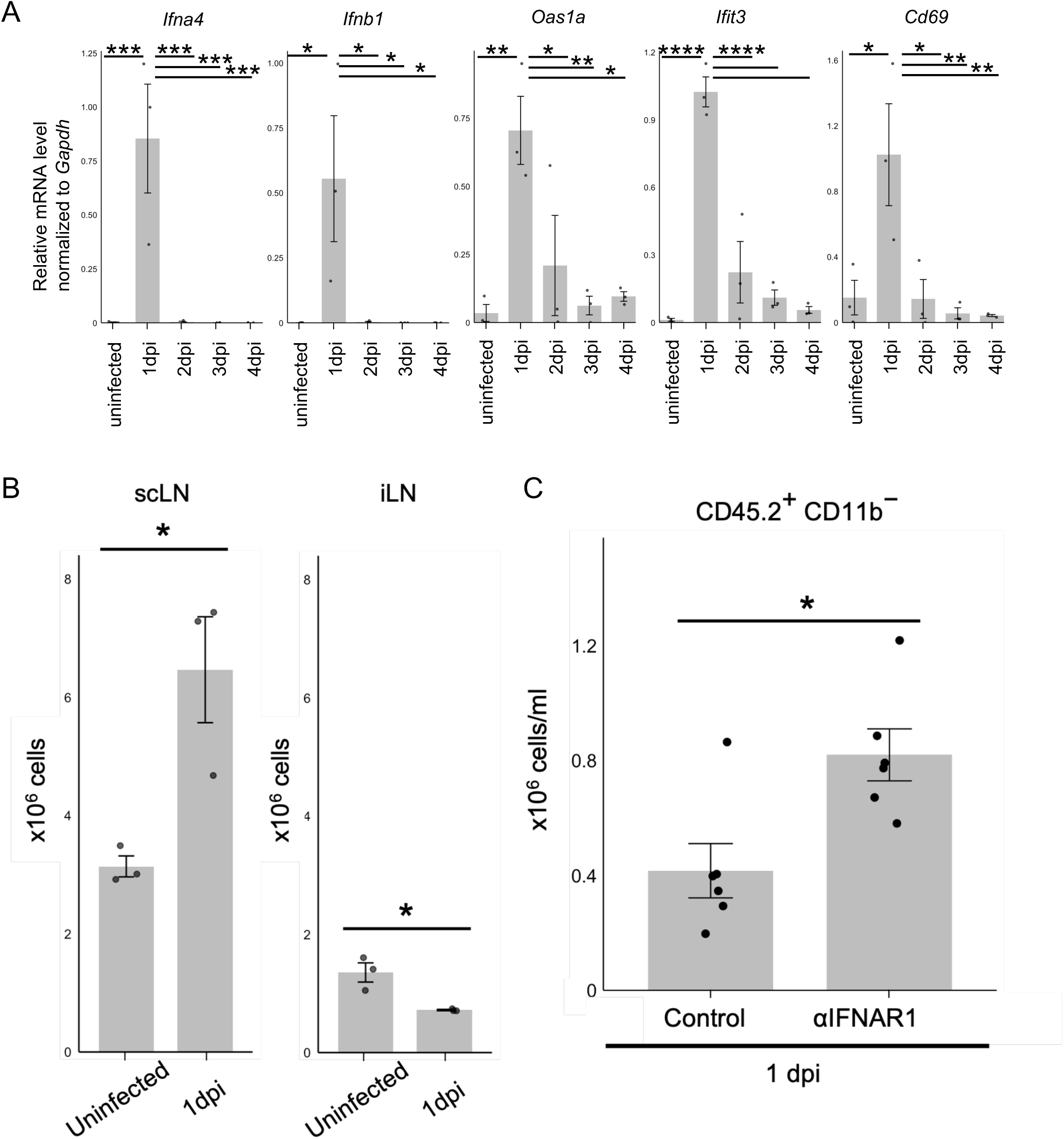
Transient type I interferon induction in draining lymph nodes mediates lymphocyte retention and systemic immune remodeling. **(A)** Relative mRNA levels of *Ifna4*, *Ifnb1*, *Oas1a*, *Ifit3*, and Cd69 in superficial cervical lymph nodes (scLNs) from uninfected and infected mice at the indicated dpi (n = 3 each). **(B)** Absolute numbers of CD45⁺CD11b⁻ lymphocytes in the scLNs and inguinal lymph nodes (iLNs) from uninfected and infected mice at 1 dpi (n = 3 each). **(C)** Absolute numbers of circulating CD45⁺CD11b⁻ lymphocytes at 1 dpi from mice treated with anti-IFNAR1 antibody or isotype IgG at 0 dpi (n = 6 each). Data are presented as mean ± SEM (A, B and C). \**P* < 0.05, \*\**P* < 0.01, \*\*\**P* <0.005, \*\*\*\**P* <0.001. Statistical significance was determined using one-way ANOVA with Tukey’s post-hoc test (A) or an unpaired, two-tailed Student’s t-test (B and C).

### Temporal requirement of type I IFN signaling governs a critical window of survival following VSV infection

Given that type I IFN is essential for antiviral defense, we hypothesized that divergent survival outcomes after intranasal VSV challenge might stem from differences in the initial type I IFN response. To test this, mice were retrospectively classified into survival and lethal groups (Fig. 3A), and plasma IFN-β levels were measured at 1 dpi (Fig. 3B). Strikingly, IFN-β levels were significantly higher in the survival group than in the lethal group, indicating that a robust early type I IFN response is associated with a favorable outcome. This notion is corroborated by the observation that survival group, which had higher IFN-β levels, similarly exhibited a more pronounced reduction in circulating lymphocytes at 1 dpi (Fig.3C). To directly evaluate the functional requirement for type I IFN signaling, we administered an anti-IFNAR1 antibody. Blocking type I IFN signaling markedly reduced survival when treatment was initiated at 0 dpi or 1 dpi but, importantly, had a less pronounced effect when given at 2 dpi (Fig. 3D). These findings identify a narrow and critical window immediately after infection during which type I IFN signaling is indispensable, demonstrating that the timing of this response is a key determinant of disease outcome.

**Figure 3.**
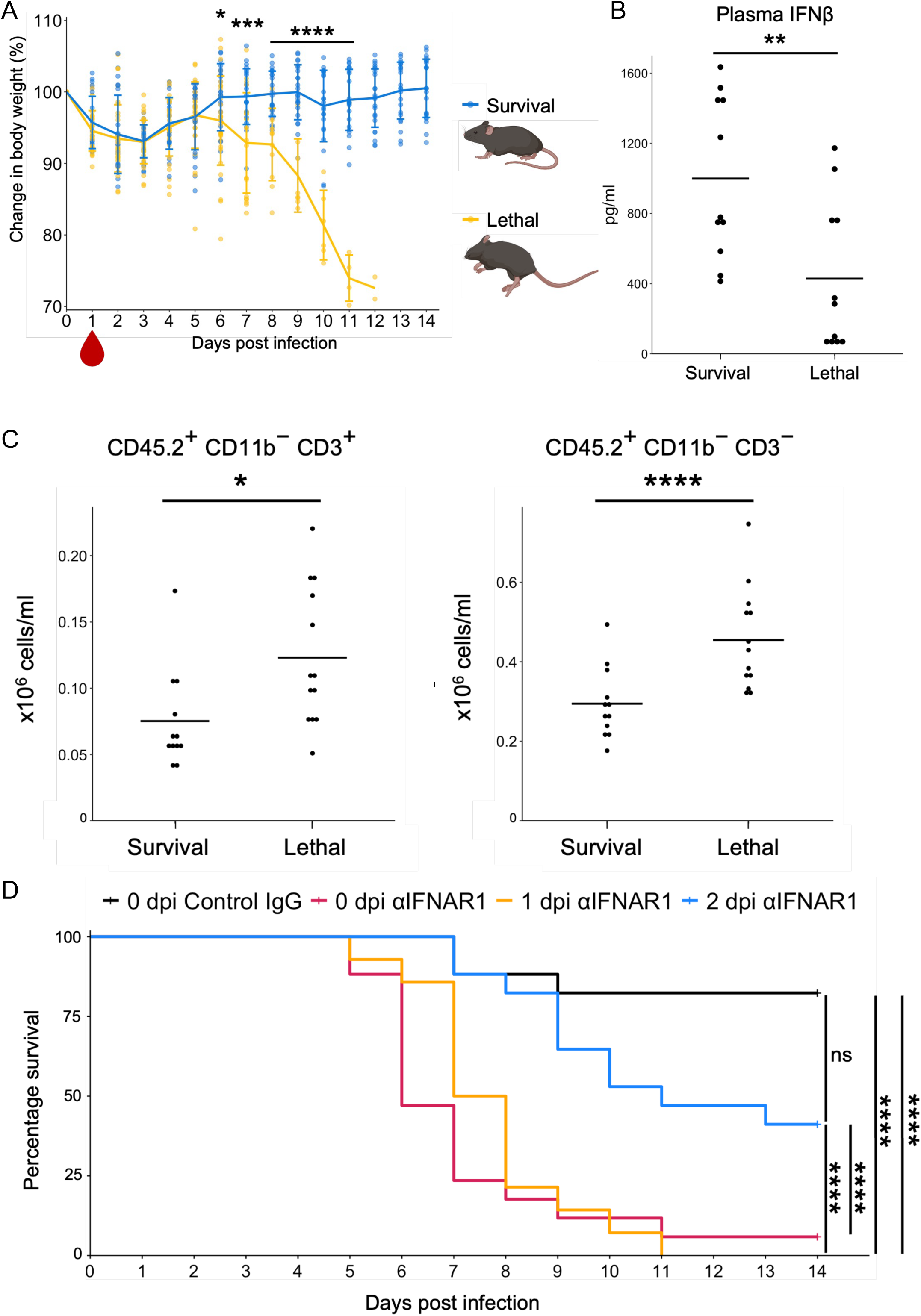
A temporally restricted type I IFN surge is essential for the survival. **(A)** Change in body weight of mice in the survival and lethal groups (n = 23 and 33, respectively). **(B)** IFNβ levels in the plasma from the survival and lethal groups (n = 11 each) at 1 dpi, measured by ELISA. **(C)** Absolute numbers of circulating CD45^+^ CD11b⁻ lymphoid cells in the survival and lethal groups at 1 dpi (n = 12 and 13, respectively). **(D)** Survival curves of mice treated with anti-IFNAR1 at 0 dpi (n = 17), 1 dpi (n = 14), or 2 dpi (n = 17), or with isotype IgG at 0 dpi (n = 17). Data in (A) are presented as mean ± SD. Data in (B and C) are shown as a dot plot where each point represents an individual mouse and the horizontal line indicates the mean. \*\**P* < 0.01. Statistical significance was determined using an unpaired, two-tailed Mann-Whitney U test (A, B and C) or the Wilcoxon (Breslow) test with Benjamini-Hochberg (BH) correction for multiple comparisons (D).

### ICAM1 expression defines an early neutrophil signature associated with survival outcome

Given that an early type I IFN surge is associated with survival, we next compared the systemic immune profiles of the survival and lethal groups to characterize the cellular response during this critical window. The most striking difference was observed in the neutrophil compartment. Specifically, a neutrophil cluster designated ISG_high_2 constituted a higher percentage of circulating cells in the survival group, while the ISG_high_1 cluster was more prevalent in the lethal group (Fig. 4A).

**Figure 4.**
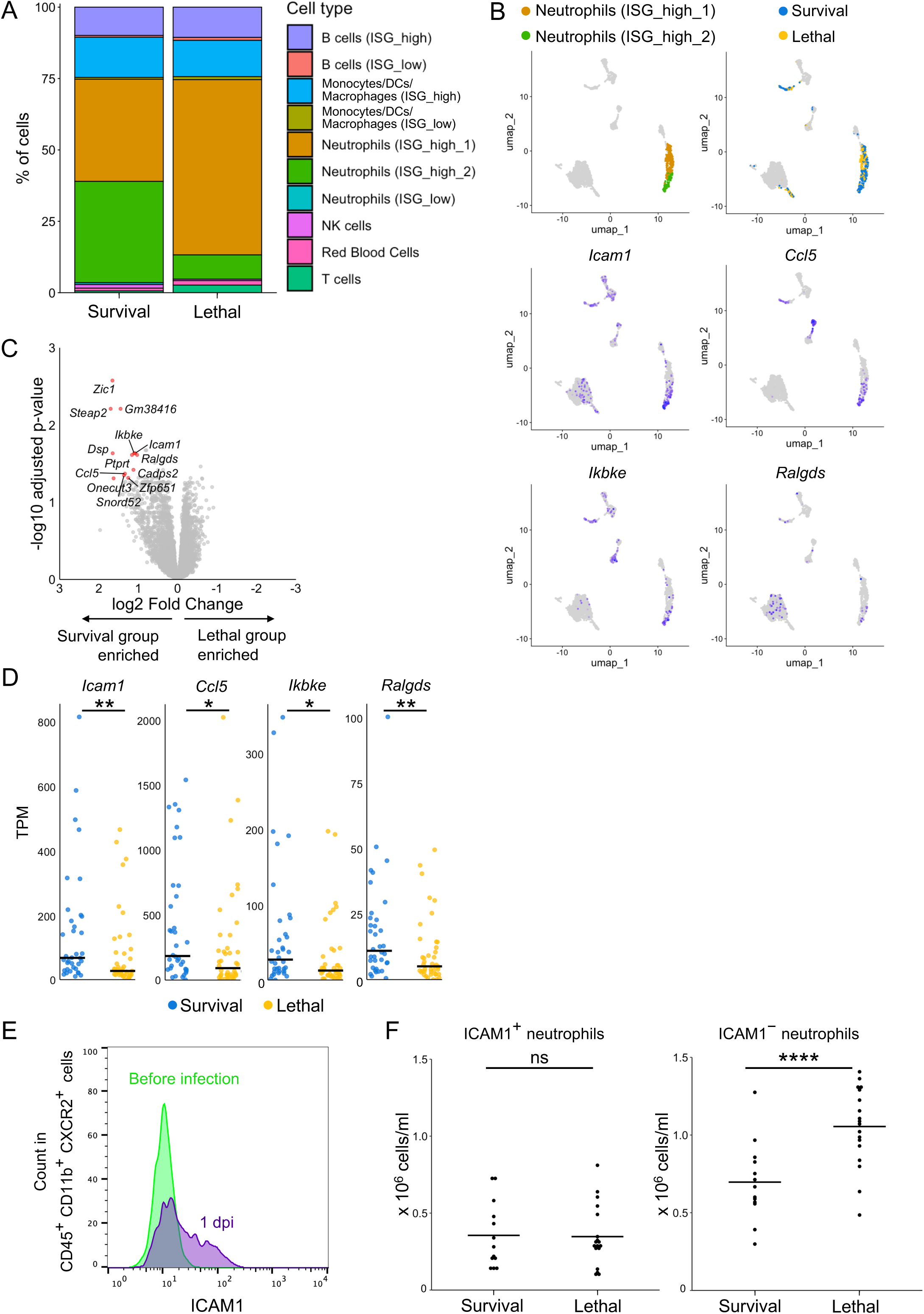
ICAM1 expression distinguish early neutrophil heterogeneity associated with survival outcome. **(A)** Percentages of each cell cluster within total circulating cells in the survival and lethal groups at 1 dpi. **(B)** UMAP and feature plots from scRNA-seq analysis, showing expression of key differentially expressed genes (DEGs) between Neutrophils (ISG_high_1) and Neutrophils (ISG_high_2) . **(C)** Volcano plot depicting DEGs in circulating neutrophils from the survival group versus the lethal group (n = 36 and 42, respectively) at 1 dpi. Genes with an absolute log2 fold change > 1 and adjusted p-value < 0.05 are highlighted in red. **(D)** Expression levels (TPM; transcripts per million) of key DEGs identified in both the bulk RNA-seq of sorted CXCR2⁺ cells and the scRNA-seq of neutrophils. **(E)** Representative histograms of ICAM1 surface expression on blood neutrophils (CD45⁺CD11b⁺CXCR2⁺) before infection and at 1 dpi. **(F)** Absolute numbers of ICAM1⁺ and ICAM1⁻ neutrophils in the blood of the survival and lethal groups (n = 13 and 18, respectively) at 1 dpi. Data in (D and F) are shown as a dot plot where each point represents an individual mouse and the horizontal line indicates the mean. ns, not significant, \**P* < 0.05, \*\**P* < 0.01, \*\*\*\**P* <0.001. Statistical significance was determined using an unpaired, two-tailed Mann-Whitney U test (D and F).

We next sought to further characterize the neutrophil subsets that correlate with survival. We compared the transcriptional differences between the survival-associated ISG_high_2 neutrophil cluster and the lethal-associated ISG_high_1 neutrophil cluster, which revealed that genes related to innate immune responses and pro-inflammatory signaling pathways were enriched in the survival-associated ISG_high_2 neutrophil cluster (Fig. S2A, B), such as *Icam1* and *Ccl5* (Fig. 4B). We also performed bulk RNA-seq on sorted CXCR2⁺ neutrophils, which corroborated the scRNA-seq findings, demonstrating that neutrophils from the survival group expressed significantly higher levels of these signature genes (Fig. 4C, D). Among these, we focused on Icam1 because it showed robust upregulation in the survival group. In addition, its cell surface expression allows detection by flow cytometry, making it a strong candidate for a prognostic predictor. Hence, we examined ICAM1 protein levels on neutrophils by flow cytometry. Surface ICAM1 expression was upregulated at 1 dpi compared to baseline (Fig. 4E, S2C). Importantly, the lethal group exhibited a significantly higher abundance of ICAM1⁻ neutrophils at 1 dpi (Fig. 4F). This suggests that the proportional differences in neutrophil clusters observed by scRNA-seq are primarily driven by the predominance of the ICAM1⁻ population in the lethal group. Collectively, these multi-platform analyses pinpoint the early polarization of circulating neutrophils, distinguished by differential ICAM1 expression, as a primary cellular correlate of survival.

### Systemic type I IFN acts on the bone marrow to induce ICAM1 expression on neutrophils

To establish a mechanistic link between the early type I IFN surge and the subsequent emergence of ICAM1⁺ neutrophils, we investigated whether type I IFN could directly regulate ICAM1 expression. Neutrophils are generated from myeloid progenitors in the bone marrow (BM) and subsequently enter the circulation^24^. Thus, type I IFN could influence ICAM1 expression on neutrophils both in the BM and in the circulation. To test these possibilities, we administered IFN-β to mice and analyzed neutrophils in both peripheral blood and BM 24 hours later. IFN-β treatment did not increase the proportion of ICAM1⁺ neutrophils in peripheral blood; however, it significantly increased the proportion of ICAM1⁺ neutrophils in the BM (Fig. S3A). These findings led us to hypothesize that infection-induced type I IFN responses act within the BM to generate an ICAM1⁺ neutrophil population, which later appears as distinct subsets in the circulation. To test the prediction that the survival group should exhibit a stronger BM type I IFN response, we retrospectively compared transcriptomic profiles of BM cells. Specifically, we performed BM biopsy at 1 dpi, which represents an early time point for the divergence of disease outcomes. We then classified the samples into survival and lethal groups based on their ultimate fate. This analysis revealed that genes expressed higher in the survival group were significantly enriched for pathways associated with IFN-β responses (Fig. S3B, C). Together, these results support a model in which mice in the survival group mount a more robust type I IFN response in the BM, leading to an increased number of ICAM1⁺ neutrophils. This early priming at the BM may underlie the differences in circulating neutrophil subsets associated with survival outcomes.

### Neutrophil Heterogeneity Defined by ICAM1 Correlates with Transcriptional and Functional Polarization

To gain insights into whether these two neutrophil populations exhibit distinct antiviral properties, we first compared the transcriptomic differences between two neutrophil populations. We performed RNA-seq on sorted ICAM1⁺ and ICAM1⁻ neutrophils, which revealed that ICAM1⁺ neutrophils were enriched for genes related to innate immune responses and pro-inflammatory signaling pathways (Fig. 5A, B). This transcriptomic profile mirrors that of the survival-associated ISG_high_2 cluster compared to the lethal-associated ISG_high_1 cluster in scRNA-seq analysis (Fig. S2A, B). In contrast, genes highly expressed in ICAM1⁻ neutrophils did not reveal clear enrichment in any specific pathways (Fig. S4A). Highly inflammatory transcriptional signature of ICAM1^+^ neutrophils is reminiscent of the N1 polarization of neutrophils in tumor microenvironment^25^. Indeed, ICAM1⁺ neutrophils were significantly enriched for a pro-inflammatory N1-like signature compared to ICAM1⁻ neutrophils (Fig. 5C), consistent with higher expression of N1 signature genes in the survival-associated ISG_high_2 cluster relative to the lethal-associated ISG_high_1 cluster (Fig.S4B). These results suggest that survival-associated neutrophils resemble the transcriptome of functionally competent N1 phenotype described in tumor microenvironments, in the context of acute viral infection.

**Figure 5.**
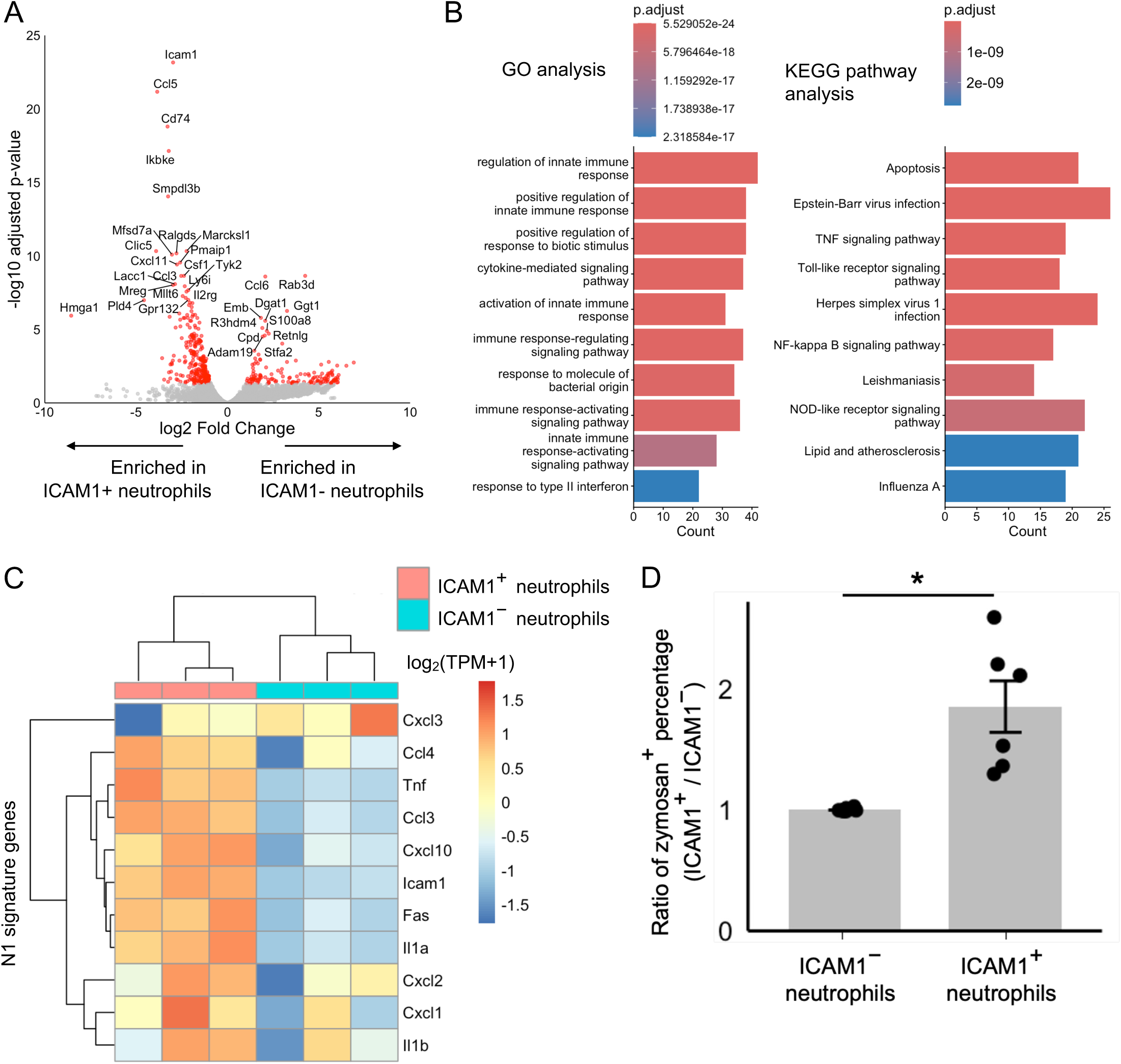
ICAM1 expression distinguishes transcriptionally and functionally distinct neutrophil subsets associated with survival outcome. **(A)** Volcano plot depicting DEGs in ICAM1⁺ neutrophils compared to ICAM1⁻ neutrophils from sorted blood samples (n = 3 per group) at 1 dpi. Genes with an absolute log2 fold change > 1 and adjusted p-value < 0.05 are highlighted in red. **(B)** Gene Ontology (GO) and KEGG pathway analysis of 293 genes more highly expressed in ICAM1⁺ neutrophils compared to ICAM1^-^ neutrophils (adjusted p-value < 0.05, log2FoldChange < -1). P-values were determined by a hypergeometric test with BH correction. **(C)** Heatmap showing the expression of N1 signature genes in sorted ICAM1⁺ and ICAM1⁻ neutrophils. **(D)** In vivo phagocytic activity of ICAM1⁺ versus ICAM1⁻ neutrophils, measured as the percentage of cells that engulfed fluorescently labeled zymosan particles. For each mouse, the value for the ICAM1⁺ population was normalized to that of the ICAM1⁻ counterpart (set as 1) (n = 6). Data in (D) are presented as mean ± SE. \**P* < 0.05. Statistical significance was determined using a one-sample t test (D).

Next, we asked whether the transcriptional polarization we observed is coupled with functional divergence in our model. ICAM1^+^ neutrophils have been reported to be a potent cell population, exhibiting enhanced effector functions such as greater phagocytic activity in a murine model of endotoxemia^26^. To determine if this functional advantage holds true in the context of viral infection, we assessed their phagocytotic function. Indeed, circulating ICAM1^+^ neutrophils from VSV-infected mice at 1dpi showed significantly higher phagocytic activity than their ICAM1⁻ counterparts (Fig. 5D). This indicates that ICAM1⁺ neutrophils represent a functionally more potent subset during the viral infection. Collectively, these findings reveal that ICAM1 expression defines a transcriptional and functional polarization of neutrophils, distinguishing a potent, pro-inflammatory subset from a distinct, less-activated population.

## DISCUSSION

A fundamental challenge in infectious disease research is elucidating the precise early events that determine the ultimate clinical outcome. This is particularly difficult because most studies, especially in humans, rely on samples collected well after the initial infection or the onset of symptoms. By capturing the immune profile during the initial phase of infection and retrospectively correlating it with survival, our study not only provides a detailed characterization of systemic immune responses—such as type I IFN-mediated lymphocyte migration and myeloid activation—at an exceptionally early stage (24 hours post-infection) but also directly links these events to the host’s ultimate fate. This early divergence in the peripheral circulation is particularly striking, as it indicates that a prognostic systemic immune signature emerges prior to the development of major cellular changes at the olfactory bulb, the primary site of intranasal VSV infection^19^. Thus, while the critical host-pathogen interactions ultimately unfold within the infection site, our findings suggest that the trajectory towards survival or death is reflected in—and likely shaped by—systemic immune events established far earlier than previously appreciated.

An intuitive explanation for such variability could be minor differences in the initial infectious dose. However, our data do not support this, as the survival rate was largely independent of the viral load within the tested range (Fig. S5A), indicating that viral dose is not the primary determinant in our model. Similarly, we considered pre-existing host-intrinsic factors known to influence infection outcomes, such as basal type I IFN levels^27^ or stress hormones^28,29^; yet, we found no significant pre-infection differences in these variables between the prospective survival and lethal groups (Fig. S5B, C). The exclusion of these potential confounders suggests that the divergence in survival is governed by previously unappreciated variability that cannot be explained by viral load or basal physiological states, thereby positioning our model as a powerful platform to identify novel regulators of host response heterogeneity.

Our study also presents a methodological advance in overcoming the logistical and technical hurdles of retrospective transcriptomic analysis in high-containment viral models. A major constraint in retrospective designs is that samples must be archived until the final prognosis is settled, precluding the immediate processing of fresh cells. To overcome this challenge, we combined a cell stabilization reagent with a cryopreservation solution, enabling the stable long-term storage of samples (see detailed methods). This approach provided a critical logistical benefit, allowing us to select and analyze samples retrospectively based on the definitive survival outcome. Furthermore, for RNA-seq analysis following FACS, viral inactivation via fixation is often mandatory but typically compromises RNA extraction efficiency. We applied a DSP fixation method^30^, which we confirmed successfully inactivated the virus while effectively preserving the integrity of labile RNA.

These optimized protocols were also instrumental in combating a significant technical challenge in the transcriptomic profiling of circulating immune cells: the instability of neutrophils. Neutrophils are highly susceptible to ex vivo apoptosis and contain low amount of RNA and abundant intracellular RNases, making their RNA notoriously difficult to analyze^31^. Consequently, many transcriptomic studies of blood focus on the more stable peripheral blood mononuclear cell (PBMC) fraction, often overlooking the neutrophil compartment. Our methodology was therefore essential for robustly profiling these fragile populations and ultimately identifying the ICAM1-defined neutrophil subset as a key predictor of survival.

The observation that ICAM1 expression on neutrophils is robustly detectable in peripheral blood underscores its high translational potential as a biomarker for predicting disease severity at the earliest stages of infection, as blood samples can be routinely collected in clinical settings. Notably, ICAM1 upregulation on neutrophils has been reported under inflammatory conditions in humans^32,33,34^. A critical next step is to validate these immune signatures as prognostic biomarkers in human viral infections, which could open new avenues for host-directed therapies aimed at shaping a protective immune response.

Our findings highlight a critical connection between the early type I IFN response, the subsequent shaping of neutrophil phenotypes, and a protective outcome. This is particularly noteworthy because sustained type I IFN signaling and pro-inflammatory neutrophils are often associated with immunopathology and severe outcomes in the later stages of viral infections^3,5, 35,36,37,38^. This contrast underscores the importance of timing; our data suggest that the beneficial role of the type I IFN-neutrophil axis may be strictly restricted to the early phase of infection. We further propose that type I IFN modulates neutrophil heterogeneity during their development in the bone marrow (BM). Given that elevated ICAM1 expression is associated with stress myelopoiesis^39,40^, the ICAM1⁺ population may represent a cohort of newly generated cells emerging from enhanced myelopoiesis triggered by the initial antiviral response.

The precise mechanisms by which this neutrophil heterogeneity dictates the survival outcome remain an important area for future investigation. One possibility is that the ICAM1⁺ population directly confers protection. ICAM1 expression on neutrophils is associated with enhanced anti-pathogenic activities^26, 41^, including phagocytosis, suggesting these cells have a greater capacity to eliminate the virus. Furthermore, as a canonical adhesion molecule, ICAM1 plays a fundamental role in mediating cell-cell interactions. Since ICAM1⁺ neutrophils have been reported to modulate lymphocyte activity via direct contact^42,43^, this population may further contribute to survival by shaping an optimal adaptive immune response. Conversely, the role of the ICAM1⁻ population also warrants critical consideration. While our data indicate that these cells do not show specific pathway enrichment or fully recapitulate a canonical immunosuppressive N2 phenotype at the transcriptomic level (Fig. S6A, B), their predominance in the lethal group—which exhibits a weaker type I IFN response—raises the possibility that they are not simply less-activated bystanders. Rather, this population, emerging in the context of a suboptimal type I IFN response, may actively contribute to a detrimental outcome, perhaps through ineffective viral control or other uncharacterized pathogenic functions. Therefore, future studies should investigate both the protective functions of ICAM1⁺ neutrophils and the potentially harmful roles of their ICAM1⁻ counterparts.

Beyond these potential applications in both clinical and basic research methodology, our work establishes a conceptual framework: treating innate biological variability not as experimental noise but as a discovery tool capable of revealing critical early checkpoints in pathogenesis that are otherwise hidden. This approach opens the door to a new understanding of individuality in disease, where the outcome is shaped not only by the genetic blueprint but also by the emergent dynamics of the initial immune response.

## Method

### Mice

C57BL/6 male mice (8-10 weeks old) were purchased from CLEA Japan (MGI:3055581) and used for all experiments unless otherwise noted.

Euthanasia was performed through isoflurane overdose followed by cervical dislocation. All mice were maintained and handled according to protocols approved by the Animal Care and Use Committee of the University of Tokyo and Hokkaido University.

### Infections

Vesicular stomatitis virus (VSV) serotype New Jersey was inoculated intranasally in a 50 µl PBS volume under 5% isoflurane anesthesia. The viral dose was 1.0 × 10^8^ PFU for scRNA-seq and 1.0 × 10^7^ PFU for all other experiments unless otherwise specified. Mice that lost more than 30% of their initial body weight or survived longer than 14 days post-infection were euthanized in accordance with humane endpoint guidelines.

### Blood and tissue sampling

150 µl of blood was collected from the tail artery into a tube containing 100 µl of 50 mM EDTA-2K in PBS. The red blood cells were lysed with ACK lysing buffer (Thermo, A1049201).

For the tissue dissection, mice were anesthetized with 200 µl of combination anesthetic (containing 75 µg/ml Medetomidine Hydrochloride, 400 µg/ml Midazolam, and 500 µg/ml Butorphanol Tartrate in saline). After perfusion with 10 ml of PBS, superficial cervical lymph nodes (scLN) and inguinal lymph nodes (iLN) were collected. Cells were collected by mincing the lymph nodes through 35μm nylon mesh. Bone marrow components were sampled by flushing the femur with 10ml 1% Albumin, from Bovine Serum (BSA), Fatty Acid Free (Fujifilm, 011-15144) in HBSS(+) with 23G syringe.

For the biopsy of femur bone marrow cells, mice anesthetized with 200 µl of combination anesthetic and cells were aspirated with a syringe 27G needle inserted directly from knee into the femur. Mice were treated with 75 µg/ml Atipamezole Hydrochloride solution (200 µl) to antagonize the anesthesia.

### ELISA

Whole blood samples in EDTA solution were centrifuged for 10 min at 1500 × g/4°C. The supernatants were collected as plasma samples and stored at -80°C. Concentrations of IFNβ(PBL Assay Science, No.42400-1) and corticosterone (Enzo Life Sciences, ADI-900-097) were determined by ELISA according to the manufacturer’s instructions.

### Cytokine stimulations

For in vivo IFN-β stimulation, recombinant IFN-β1 (carrier-free) (PBL assay science, 12401, or Biolegend, 581302) diluted with 0.1% BSA/PBS was used. Uninfected mice received intravenous (i.v.) injection of 2.78 × 10^4^ units IFN-β in 100μl solution. 0.1% BSA/PBS was injected for control. The blood and the bone marrow was sampled after 24 hours. Samples were processed for flowcytometry analysis.

### Anti-IFNAR1 antibody treatment in vivo to inhibit type I IFN responses

Mice were treated with Anti-mouse IFNAR-1-InVivo (Selleck, A2121, 500μg in 200μl PBS i.p. at the time of infection, 1 dpi or 2 dpi) or Mouse IgG1 isotype control-InVivo (Selleck, A2106).

### Zymosan administration in vivo to evaluate phagocytotic activity

Mice received injection of 100μg Alexa Fluor 488 conjugated Zymosan A BioParticles (Invitrogen, Z23373) in 100μl PBS at day1 after infection. The blood was sampled after 15 minutes and processed for flowcytometry analysis.

### Flow cytometry

Collected cells were incubated at 4°C for 15 min with 4% paraformaldehyde (PFA) (Sigma-Aldrich, 441244). After centrifugation for 5 min at 800 × g/4°C, the cell suspensions were washed and resuspended in 3% BSA/PBS. The suspensions were incubated at 4°C for 30 min for surface staining. The list of flow antibodies is provided in key resources table. The samples were washed and resuspended in 0.2% BSA/PBS and filtered through a 35 µm mesh filter (Falcon). Flowcytometry data were acquired with FACS Aria III, IIIu (BD Biosciences) or CytoFLEX SRT (Beckman Coulter). Data was analyzed using FlowJo version 10 software. The following antibodies were used: PerCP/Cyanine5.5 anti-mouse CD45.2 clone 104 (1:200, Biolegend, 109828), APC/Cyanine7 anti-mouse CD11b clone M1/70 (1:200, Biolegend, 101226), Purified anti-mouse CD16/32 clone 93 (1:50, Biolegend, 101302), PE anti-mouse CD182(CXCR2) clone SA044G4 (1:200, Biolegend, 149304), APC anti-mouse CD54(ICAM1) clone YN1/1.7.4 (1:300, Biolegend, 116129).

### Fluorescence-activated cell sorting (FACS)

Prior to sorting ICAM1^+^ and ICAM1^-^ neutrophils, cells were fixed with dithio-bis-succinimidyl propionate (DSP) (Thermo Scientfic, 22586) as previously described^44^. This method inactivates the virus while preserving RNA integrity for downstream analysis. These fixed cells were then subjected to the staining protocol described above and were collected with using FACS Aria III or IIIu (BD Biosciences). Gating for sorting was applied using FACSDiva version 6.

### Magnetic Activated Cell Sorting (MACS)

For neutrophil isolation for bulk RNA-seq, blood-derived cells were incubated at 4°C for 15 min firstly with anti-CXCR2 antibody (1:50, clone SA044G4, BioLegend, 149304) and subsequently with anti-PE microbeads (1:100, Miltenyi Biotec, 130-048-801). Subsequently, cells were subjected to MACS (Miltenyi Biotec) and magnetically retained cells were eluted from the MS column and collected by centrifugation for 5 min at 400 x g/4°C.

### Single-cell RNA sequencing and analysis

Blood was sampled at 0, 1 and 5 dpi as described in the previous section. Cells were fixed with CellCover (Anacyte, 800-125) at 4℃ for 15min to preserve RNA. Cells were then resuspended in CELLBANKER 1 (Nippon Zenyaku Kogyo Co., Ltd., 11910) and stored at -80℃. After the final survival outcome was observed, samples from the survival (n=2) and lethal (n=2) groups were processed for subsequent library preparation following mRNA Whole Transcriptome Analysis (WTA) and Sample Tag Library Preparation Protocol from BD Rhapsody™ Single-Cell Analysis System with BD Mouse Immune Single-Cell Multiplexing Kit (BD, 633793). Sequencing was carried out on an HiSeq X Ten platform (Macrogen Japan) with paired-end 150 bp reads. Raw sequencing data were processed and analyzed using the BD Rhapsody™ WTA Pipeline, executed on the Seven Bridges Platform (Seven Bridges Genomics, Inc., https://www.sevenbridges.com). The pipeline included read alignment, UMI-based de-duplication, and generation of gene expression matrices based on BD’s reference transcriptome. In total, 1,710 single cells were sequenced: 166 and 208 cells at 0 dpi, 313 and 189 cells at 1 dpi, 515 and 319 cells at 5 dpi, from the survival and lethal groups, respectively. Cells expressing < 200 features, > 2,500 features, or > 25% mitochondrial genes were excluded from the analysis. Dimensionality reduction and clustering analysis were performed using Seurat v5.3.0^45^. The following cells were annotated using conventional cell markers: B cells (*Cd79a, Cd19, Ms4a*1), neutrophils (*Cxcr2, S100a8, S100a9*), Monocytes/DCs/Macrophages (*Cx3cr1, Csf1r, Adgre1*), T cells (*Cd3e*), NK cells (*Klrb1c, Nkg7*), and red blood cells (*Hba-a2*).

### RNA extraction, reverse transcription, and qPCR

Total RNA was isolated from isolated cells or tissues of mice with the use of RNAiso Plus reagent (Takara, 9109) and ReverTra Ace qPCR Master Mix (Toyobo, FSQ301) following manufacturer’s protocols. The resulting cDNA was subjected to real-time PCR analysis in a LightCycler 96 instrument (Roche) with KAPA SYBR Fast qPCRKit (Kapa Biosystems). The amount of target mRNA was normalized by that of *Gapdh* mRNA.

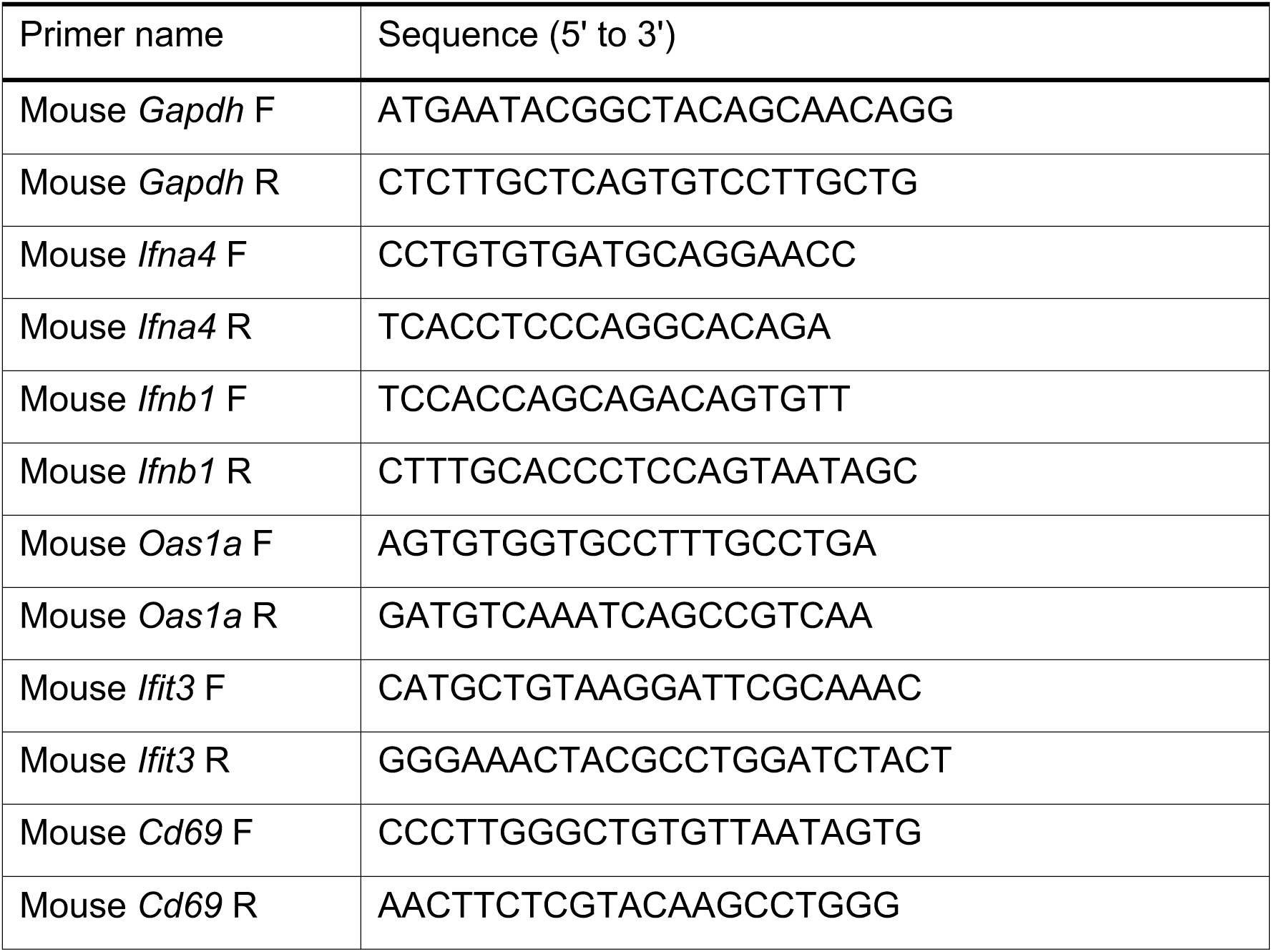

### RNA sequencing analysis

Total RNA was isolated from the cells for library construction. RNA libraries were prepared using SMART-Seq Stranded Kit (Takara) following manufacturer’s protocols. Sequencing was carried out on an Illumina NextSeq 2000 platform with paired-end 36 bp reads or on a DNBSEQ-G400RS platform with 100 bp single-end reads. For DNBSEQ-G400RS sequencing, adaptor sequences were converted into the DNBSEQ format using the MGIEasy Universal Library Conversion Kit (App-A) (MGI). Sequences were mapped to the mouse reference genome (mm10) with the use of STAR v 2.7.10a^46^. Raw read counts of each gene were calculated using featureCounts v2.0.^47^. Adjusted p-values and fold changes used to identify differentially expressed genes (DEGs) were calculated using the DESeq2 (v1.44.0) R package^48^. Gene Ontology (GO) enrichment analysis for biological processes (BP) and Kyoto Encyclopedia of Genes and Genomes (KEGG) pathway enrichment analysis were conducted using the enrichGO and enrichKEGG functions in the clusterProfiler (v4.12.6) R package ^49^, with org.Mm.eg.db and mouse genes (organism code: “mmu”) as annotation databases, respectively. Only terms and pathways with adjusted p-values < 0.05 were considered significant. Plots were generated using ggplot within the ggplot2 package 3.5.2^50^. The N1 and N2 signature genes shown in Fig. 5C and S6A were selected based on previous studies^51,52,53,54,55^.

### Statistical analysis and data availability

All data were plotted and analyzed using R software (v.4.4.0)^56^ driving RStudio (v.2024.4.2.764). Data are presented as mean ± s.e.m. including the individual values where possible, and n represents the number of individual animals (biological replicates) unless otherwise noted. All flow cytometry images shown are representative of at least three independent experiments repeated with similar results. Means between two groups were compared with an unpaired or a paired two-tailed t tests, or an unpaired two-tailed Mann-Whitney U test. Means among three or more groups were compared with one-way ANOVA with Tukey’s post-hoc test. When values were normalized to one condition (e.g., ICAM1⁻), a one-sample two-tailed t test was used to compare the normalized values to the hypothetical value of 1. Survival curves were generated using the Kaplan–Meier method and compared using the Wilcoxon (Breslow) test with BH correction for multiple comparisons. scRNA-seq and RNA-seq data have been deposited in the Gene Expression Omnibus (GEO) under accession number GSE313167, a SuperSeries that comprises GSE313019, GSE313732, GSE313164, and GSE313165. All data and codes used for data processing and statistical analysis are provided with this paper as supplemental files.

### Data availability

scRNA-seq data and RNA-seq data are deposited in the GEO: GSE313167 and are publicly available as of the date of publication. The codes used for transcriptome analyses are provided with this paper as supplemental files.

## Acknowledgements

All members of Laboratory of Molecular Biology (Graduate School of Pharmaceutical Sciences, The University of Tokyo) and Laboratory of Molecular Cell Biology (Institute for Genetic Medicine, Hokkaido University) community for feedback throughout this project.

R.S. was supported by Ministry of Education, Culture, Sports, Science, and Technology of Japan (MEXT) JP23KJ0674.

A.S. was supported by MEXT JP22KJ1176.

H.S. was supported by Japan Agency for Medical Research and Development (AMED) JP223fa627001 (UTOPIA Young Researcher Development Program).

Y.G. was supported by MEXT JP22H00431, MEXT JP22H04925(PAGS), MEXT JP24H02322, MEXT JP18gm0610013, MEXT JP24gm1310004.

T.O. was supported by MEXT JP24K02172, MEXT J24K22007, FORST program of the Japan Science and Technology Agency JPMJFR204Q, Takano Life Science Research Foundation, Chemo-Sero-Therapeutic Research Institute, Mitsubishi Foundation, Naito Foundation, Secom Foundation, and Astellas Foundation for Research on Metabolic Disorders.

## Author contributions

Conceptualization: R.S. and T.O.

Methodology: R.S., H.S.

Software: R.S., H.S.

Formal Analysis: R.S., H.S.

Investigation: R.S., A.S.

Writing – Original Draft: R.S. and T.O.

Writing – Review & Editing: All authors.

Supervision: Y.G., T.O.

Project Administration: T.O.

Funding Acquisition: H.S., Y.G., T.O.

---

**Figure S1.**
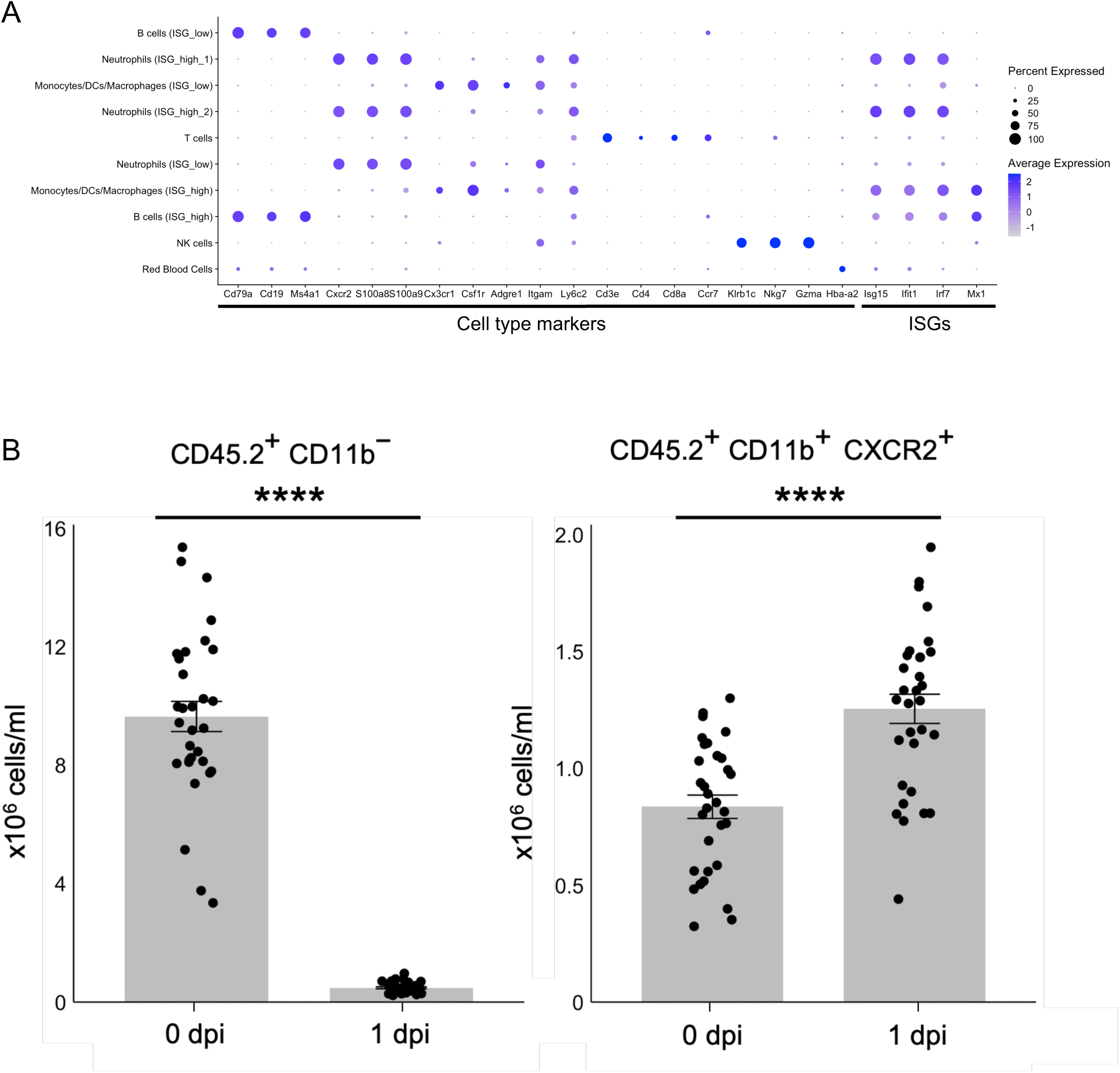
Expression profiles of canonical lineage markers and interferon-stimulated genes (ISGs) across all cell clusters. **(A)** Dot plot showing the expression of canonical marker genes for cell type identification and key ISGs across all cell clusters identified by scRNA-seq. **(B)** Absolute numbers of circulating CD45⁺CD11b⁻ lymphocytes and CD45⁺CD11b^+^CXCR2^+^ neutrophils at 0 and 1 dpi (n = 31). Data in (B) are presented as mean ± SEM. \*\*\*\**P* <0.001. Statistical significance was determined using a paired, two-tailed Student’s t-test (B).

**Figure S2.**
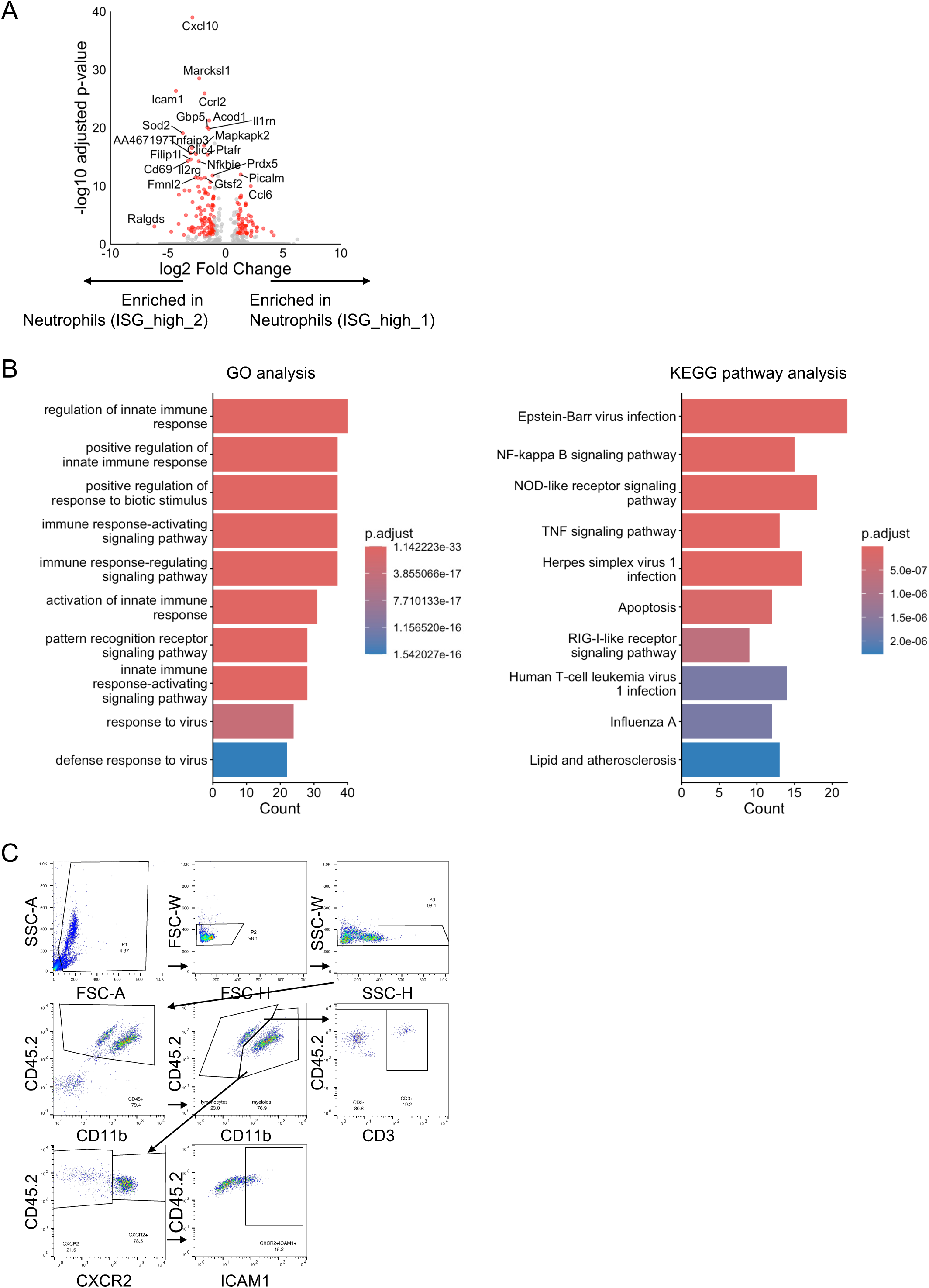
Distinct gene expression profiles between ISG-high neutrophil subsets. **(A)** Volcano plot depicting DEGs between the scRNA-seq-defined Neutrophils (ISG_high_2) cluster and Neutrophils (ISG_high_1). Genes with an absolute log2 fold change > 1 and adjusted p-value < 0.05 are highlighted in red. (n = 3 mice) **(B)** Gene Ontology (GO) and KEGG pathway analysis of 152 genes more highly expressed in the scRNA-seq-defined Neutrophils (ISG_high_2) cluster compared to the Neutrophils (ISG_high_1) cluster (adjusted p-value < 0.05, log2FoldChange < - 1). The top 10 enriched terms are shown. **(C)** Representative flow cytometry gating strategy for the identification of major immune cell populations in the blood

**Figure S3.**
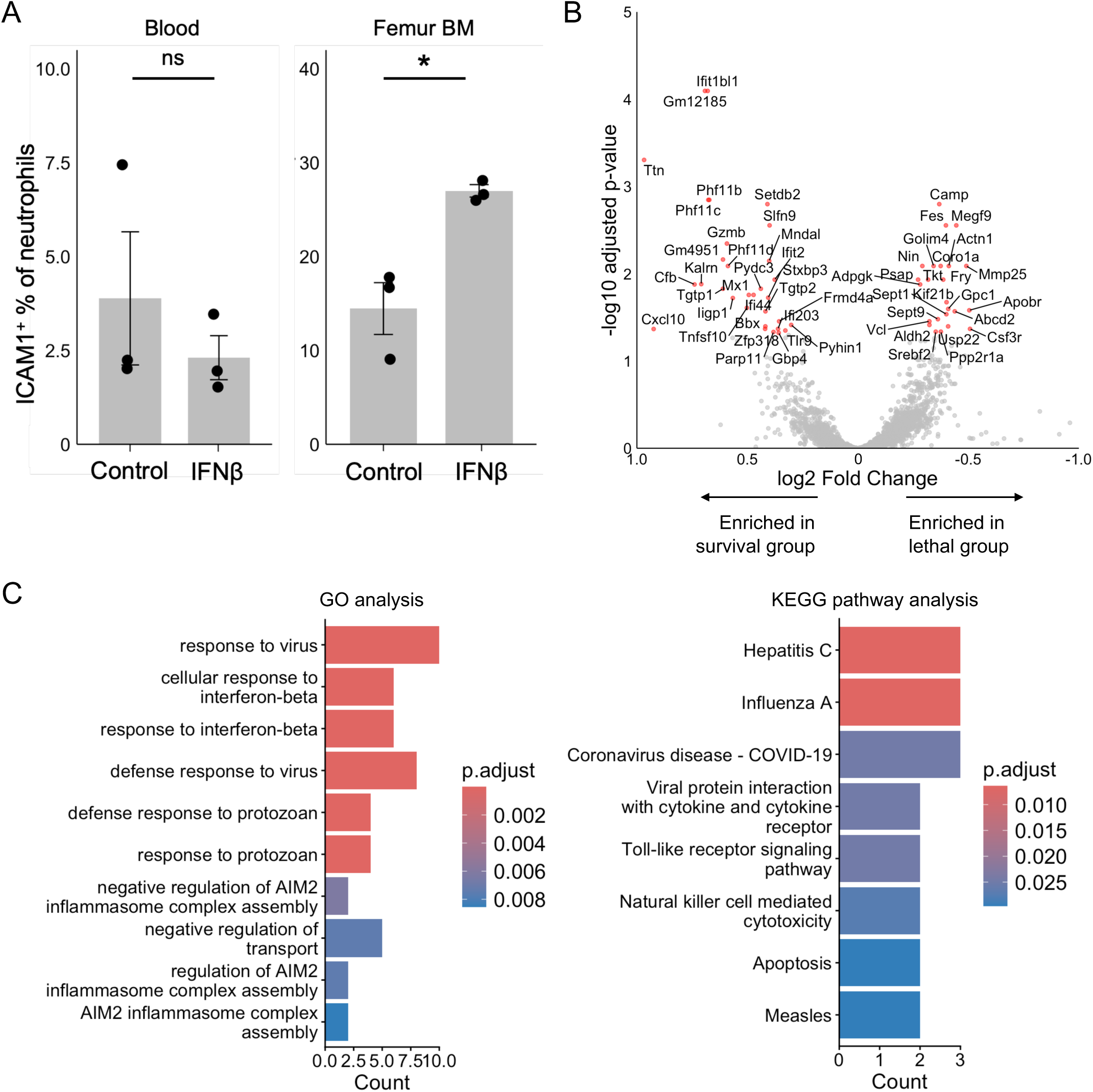
IFNβ-induced neutrophil heterogeneity not in the blood but in the peripheral bone marrow (BM), and the survived group expressed higher levels of IFNβ signaling related genes in the bone marrow at 1 dpi. **(A)** Percentages of ICAM1^+^ cells within CXCR2^+^ neutrophils in the blood and the the femur BM at 24 hours after IFNβ administration (n = 3 mice). **(B)** Volcano plot depicting DEGs in CXCR2^+^ neutrophils in the femur BM from the survived versus the lethal group (n = 6 each) at 1 dpi. Genes with an adjusted p-value < 0.05 are highlighted in red. **(C)** Gene Ontology (GO) and KEGG pathway analysis of 31 genes more highly expressed in the femur BM from the survived group compared to the lethal group (adjusted p-value < 0.05, log2FoldChange < 0). Data in (A) are shown as a dot plot where each point represents an individual mouse and the horizontal line indicates the mean. NS, not significant. \**P* < 0.05. Statistical significance was determined using an unpaired, two-tailed Student’s t-test (A).

**Figure S4.**
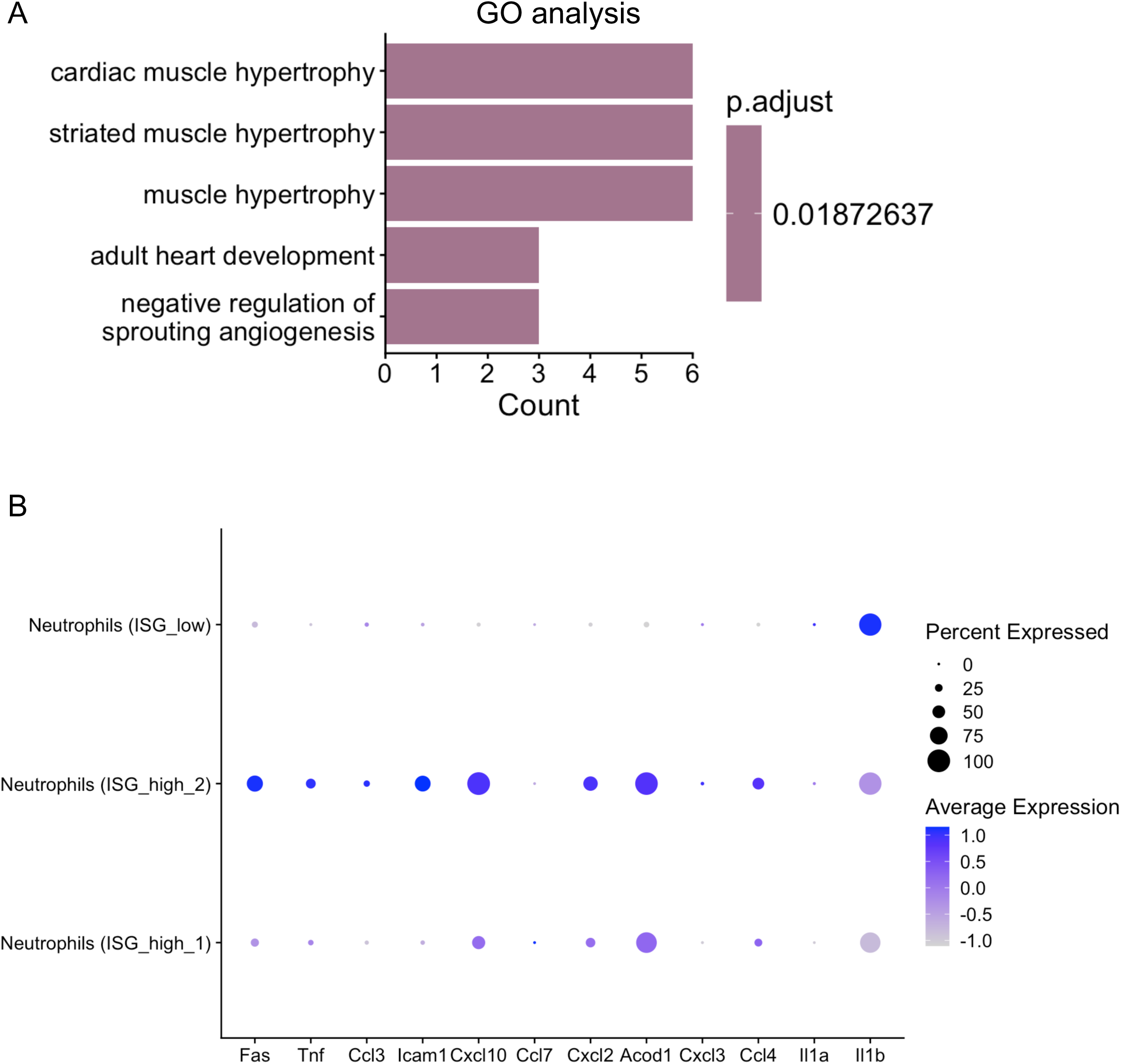
Additional characterization of circulating neutrophil subsets. **(A)** Gene Ontology (GO) and KEGG pathway analysis of 149 genes more highly expressed in ICAM1^-^ neutrophils compared to ICAM1^+^ neutrophils (adjusted p-value < 0.05, log2FoldChange > 1). No enriched pathway was found in KEGG pathway analysis with this gene sets. **(B)** Dot plot from scRNA-seq data showing expression levels of N1 signature genes within each identified cell cluster.

**Figure S5.**
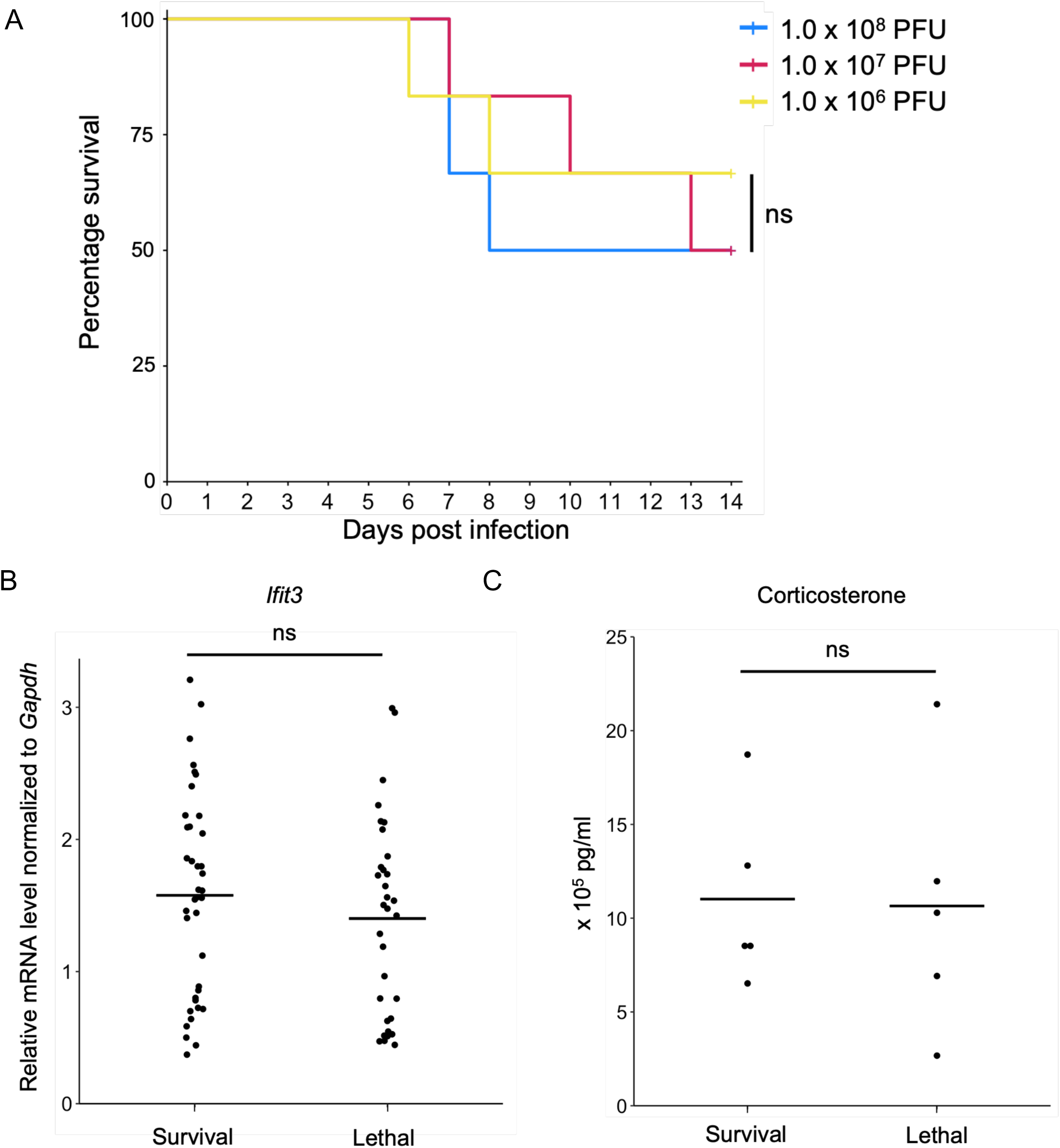
No clear association was found between viral dose, baseline ISG, or corticosterone and survival outcome. **(A)** Survival curves of mice infected with the indicated doses of VSV (n = 6 each). **(B)** Relative mRNA levels of *Ifit3* from blood circulating cells of survival and lethal groups before infection (n = 38 and n = 32, respectively). **(C)** Corticosterone levels in the plasma of the survival and lethal groups before infection (n = 5 each). Data in (B and C) are shown as a dot plot where each point represents an individual mouse and the horizontal line indicates the mean. ns, not significant. Statistical significance was determined using the Wilcoxon (Breslow) test with BH correction for multiple comparisons (A) or an unpaired, two-tailed Student’s t-test (B and C).

**Figure S6.**
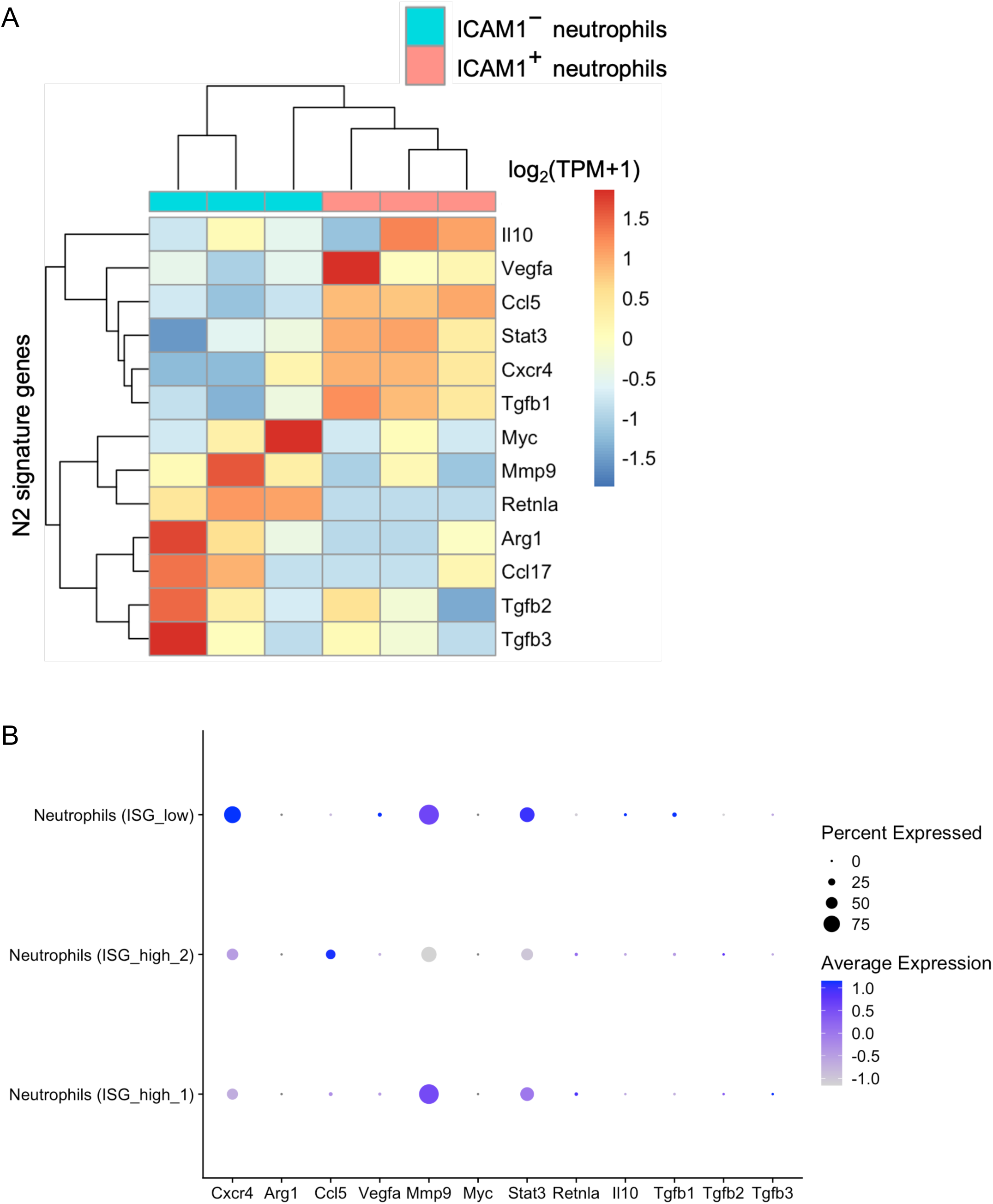
N2 signature genes are not specifically enriched in ICAM1⁻ neutrophils or the lethal group-associated ISG_high_1 cluster. **(A)** Heatmap showing the expression of and N2 signature genes in sorted ICAM1⁻ and ICAM1⁺ neutrophils. **(B)** Dot plot from scRNA-seq data showing expression levels of N2 signature genes within each identified cell cluster.

## References

1 Rouse, Barry T., and Sharvan Sehrawat. “Immunity and immunopathology to viruses: what decides the outcome?.” Nature Reviews Immunology 10.7 (2010): 514–526.

2 Liu, Rui, et al. “Decreased T cell populations contribute to the increased severity of COVID-19.” Clinica Chimica Acta 508 (2020): 110–114.

3 Silvin, Aymeric, et al. “Elevated calprotectin and abnormal myeloid cell subsets discriminate severe from mild COVID-19.” Cell 182.6 (2020): 1401–1418.

4 Lucas, Carolina, et al. “Longitudinal analyses reveal immunological misfiring in severe COVID-19.” Nature 584.7821 (2020): 463–469.

5 Arunachalam, Prabhu S., et al. “Systems biological assessment of immunity to mild versus severe COVID-19 infection in humans.” Science 369.6508 (2020): 1210–1220.

6 Schughart, Klaus, et al. “Host response to influenza infections in human blood: association of influenza severity with host genetics and transcriptomic response.” Frontiers in Immunology 15 (2024): 1385362.

7 Lindeboom, Rik GH, et al. “Human SARS-CoV-2 challenge uncovers local and systemic response dynamics.” Nature 631.8019 (2024): 189–198.

8 Hayden, Frederick G., et al. “Local and systemic cytokine responses during experimental human influenza A virus infection. Relation to symptom formation and host defense.” The Journal of clinical investigation 101.3 (1998): 643–649.

9 Huneycutt, Brandon S., et al. “Central neuropathogenesis of vesicular stomatitis virus infection of immunodeficient mice.” Journal of virology 67.11 (1993): 6698–6706.

10 Thomsen, Allan Randrup, et al. “Cooperation of B cells and T cells is required for survival of mice infected with vesicular stomatitis virus.” International immunology 9.11 (1997): 1757–1766.

11 Akira, Shizuo, Satoshi Uematsu, and Osamu Takeuchi. “Pathogen recognition and innate immunity.” Cell 124.4 (2006): 783–801.

12 Thompson, Mikayla R., et al. “Pattern recognition receptors and the innate immune response to viral infection.” Viruses 3.6 (2011): 920.

13 De Giovanni, Marco, et al. “Spatiotemporal regulation of type I interferon expression determines the antiviral polarization of CD4+ T cells.” Nature immunology 21.3 (2020): 321–330.

14 Mifsud, Edin J., Miku Kuba, and Ian G. Barr. “Innate immune responses to influenza virus infections in the upper respiratory tract.” Viruses 13.10 (2021): 2090.

15 Stetson, Daniel B., and Ruslan Medzhitov. “Type I interferons in host defense.” Immunity 25.3 (2006): 373–381.

16 McNab, Finlay, et al. “Type I interferons in infectious disease.” Nature Reviews Immunology 15.2 (2015): 87–103.

17 Channappanavar, Rudragouda, et al. “IFN-I response timing relative to virus replication determines MERS coronavirus infection outcomes.” The Journal of clinical investigation 129.9 (2019): 3625–3639.

18 Channappanavar, Rudragouda, et al. “Dysregulated type I interferon and inflammatory monocyte-macrophage responses cause lethal pneumonia in SARS-CoV-infected mice.” Cell host & microbe 19.2 (2016): 181–193.

19 Moseman, E. Ashley, et al. “T cell engagement of cross-presenting microglia protects the brain from a nasal virus infection.” Science immunology 5.48 (2020): eabb1817.

20 Iannacone, Matteo, et al. “Subcapsular sinus macrophages prevent CNS invasion on peripheral infection with a neurotropic virus.” Nature 465.7301 (2010): 1079–1083.

21 Kamphuis, Elisabeth, et al. “Type I interferons directly regulate lymphocyte recirculation and cause transient blood lymphopenia.” Blood 108.10 (2006): 3253–3261.

22 Jin, Hokyung, et al. “Increased CSF drainage by non-invasive manipulation of cervical lymphatics.” Nature (2025): 1–13.

23 Shiow, Lawrence R., et al. “CD69 acts downstream of interferon-α/β to inhibit S1P1 and lymphocyte egress from lymphoid organs.” Nature 440.7083 (2006): 540–544.

24 Borregaard, Niels. “Neutrophils, from marrow to microbes.” Immunity 33.5 (2010): 657–670.

25 Fridlender, Zvi G., et al. “Polarization of tumor-associated neutrophil phenotype by TGF-β:“N1” versus “N2” TAN.” Cancer cell 16.3 (2009): 183–194.

26 Woodfin, Abigail, et al. “ICAM-1–expressing neutrophils exhibit enhanced effector functions in murine models of endotoxemia.” Blood, The Journal of the American Society of Hematology 127.7 (2016): 898–907.

27 Gough, Daniel J., et al. “Constitutive type I interferon modulates homeostatic balance through tonic signaling.” Immunity 36.2 (2012): 166–174.

28 Luo, Zhuo, et al. “Novel insights into stress-induced susceptibility to influenza: corticosterone impacts interferon-β responses by Mfn2-mediated ubiquitin degradation of MAVS.” Signal Transduction and Targeted Therapy 5.1 (2020): 202.

29 Gagnidze, Khatuna, et al. “Nuclear receptor REV-ERBα mediates circadian sensitivity to mortality in murine vesicular stomatitis virus-induced encephalitis.” Proceedings of the National Academy of Sciences 113.20 (2016): 5730–5735.

30 Attar, Moustafa, et al. “A practical solution for preserving single cells for RNA sequencing.” Scientific reports 8.1 (2018): 2151.

31 Hatje, Klas, et al. “Comparison of Fixed Single Cell RNA-seq Methods to Enable Transcriptome Profiling of Neutrophils in Clinical Samples.” bioRxiv (2024): 2024–08.

32 Muller Kobold, A. C., et al. “Are circulating neutrophils intravascularly activated in patients with anti-neutrophil cytoplasmic antibody (ANCA)-associated vasculitides?.” Clinical & Experimental Immunology 114.3 (1998): 491–499.

33 Wang, Shan-Ze, et al. “Shedding of L-selectin and PECAM-1 and upregulation of Mac-1 and ICAM-1 on neutrophils in RSV bronchiolitis.” American Journal of Physiology-Lung Cellular and Molecular Physiology 275.5 (1998): L983–L989.

34 Elsner, Jörn, et al. “Synthesis and surface expression of ICAM-1 in polymorphonuclear neutrophilic leukocytes in normal subiects and during inflammatory disease.” Immunobiology 193.5 (1995): 456–464.

35 Tang, Benjamin M., et al. “Neutrophils-related host factors associated with severe disease and fatality in patients with influenza infection.” Nature communications 10.1 (2019): 3422.

36 Shaath, Hibah, et al. “Single-cell transcriptome analysis highlights a role for neutrophils and inflammatory macrophages in the pathogenesis of severe COVID-19.” Cells 9.11 (2020): 2374.

37 Meizlish, Matthew L., et al. “A neutrophil activation signature predicts critical illness and mortality in COVID-19.” Blood advances 5.5 (2021): 1164–1177.

38 Levy, Yves, et al. “CD177, a specific marker of neutrophil activation, is associated with coronavirus disease 2019 severity and death.” Iscience 24.7 (2021).

39 Montaldo, Elisa, et al. “Cellular and transcriptional dynamics of human neutrophils at steady state and upon stress.” Nature immunology 23.10 (2022): 1470–1483.

40 Marini, Olivia, et al. “Mature CD10^+^ and immature CD10^−^ neutrophils present in G-CSF–treated donors display opposite effects on T cells.” Blood, The Journal of the American Society of Hematology 129.10 (2017): 1343–1356.

41 Takashi, Shuji, Yoshio Okubo, and Shiro Horie. “Contribution of CD54 to human eosinophil and neutrophil superoxide production.” Journal of Applied Physiology 91.2 (2001): 613–622.

42 Hawkins, Ryder F. Whittaker, et al. “ICAM1+ neutrophils promote chronic inflammation via ASPRV1 in B cell–dependent autoimmune encephalomyelitis.” JCI insight 2.23 (2017): e96882.

43 Pillay, Janesh, et al. “A subset of neutrophils in human systemic inflammation inhibits T cell responses through Mac-1.” The Journal of clinical investigation 122.1 (2012): 327–336.

44 Attar, Moustafa, et al. “A practical solution for preserving single cells for RNA sequencing.” Scientific reports 8.1 (2018): 2151.

45 Hao, Yuhan, et al. “Dictionary learning for integrative, multimodal and scalable single-cell analysis.” Nature biotechnology 42.2 (2024): 293–304.

46 Dobin, Alexander, et al. “STAR: ultrafast universal RNA-seq aligner.” Bioinformatics 29.1 (2013): 15–21.

47 Liao, Yang, Gordon K. Smyth, and Wei Shi. “featureCounts: an efficient general purpose program for assigning sequence reads to genomic features.” Bioinformatics 30.7 (2014): 923–930.

48 Love, Michael I., Wolfgang Huber, and Simon Anders. “Moderated estimation of fold change and dispersion for RNA-seq data with DESeq2.” Genome biology 15.12 (2014): 550.

49 Yu, Guangchuang. “Thirteen years of clusterProfiler.” The Innovation 5.6 (2024).

50 Ginestet, Cedric. “ggplot2: elegant graphics for data analysis.” (2011): 245–246.

51 Piccard, H., R. J. Muschel, and Ghislain Opdenakker. “On the dual roles and polarized phenotypes of neutrophils in tumor development and progression.” Critical reviews in oncology/hematology 82.3 (2012): 296–309.

52 Chen, Fei, et al. “Neutrophils prime a long-lived effector macrophage phenotype that mediates accelerated helminth expulsion.” Nature immunology 15.10 (2014): 938–946.

53 Shaul, Merav E., et al. “Tumor-associated neutrophils display a distinct N1 profile following TGFβ modulation: A transcriptomics analysis of pro-vs. antitumor TANs.” Oncoimmunology 5.11 (2016): e1232221.

54 Cai, Wei, et al. “All trans-retinoic acid protects against acute ischemic stroke by modulating neutrophil functions through STAT1 signaling.” Journal of neuroinflammation 16.1 (2019): 175.

55 Mihaila, Andreea C., et al. “Transcriptional profiling and functional analysis of N1/N2 neutrophils reveal an immunomodulatory effect of S100A9-blockade on the pro-inflammatory N1 subpopulation.” Frontiers in immunology 12 (2021): 708770.

56 R Core Team (2024). _R: A Language and Environment for Statistical Computing_. R Foundation for Statistical Computing, Vienna, Austria. <https://www.R-project.org/>.

